# Elevated EGR1 Binding at Enhancers in Excitatory Neurons Correlates with Neuronal Subtype-Specific Epigenetic Regulation

**DOI:** 10.1101/2024.11.21.624733

**Authors:** Liduo Yin, Xiguang Xu, Benjamin Conacher, Yu Lin, Gabriela L. Carrillo, Yupeng Cun, Michael A. Fox, Xuemei Lu, Hehuang Xie

## Abstract

Brain development and neuronal cell specification are accompanied with epigenetic changes to achieve diverse gene expression regulation. Interacting with cell-type specific epigenetic marks, transcription factors bind to different sets of cis-regulatory elements in different types of cells. Currently, it remains largely unclear how cell-type specific gene regulation is achieved for neurons. In this study, we generated epigenetic maps to perform comparative histone modification analysis between excitatory and inhibitory neurons. We found that neuronal cell-type specific histone modifications are enriched in super enhancer regions containing abundant EGR1 motifs. Further CUT&RUN data validated that more EGR1 binding sites can be detected in excitatory neurons and primarily located in enhancers. Integrative analysis revealed that EGR1 binding is strongly correlated with various epigenetic markers for open chromatin regions and associated with distinct gene pathways with neuronal subtype-specific functions. In inhibitory neurons, the majority of genomic regions hosting EGR1 binding sites become accessible at early embryonic stages. In contrast, the super enhancers in excitatory neurons hosting EGR1 binding sites gained their accessibility during postnatal stages. This study highlights the significance of transcription factor binding to enhancer regions, which may play a crucial role in establishing cell-type specific gene regulation in neurons.

## Background

Millions of neurons in the mouse brain interact with each other to achieve unique functions including locomotion control, sensation, and memory. These neurons may be classified into two large categories, excitatory and inhibitory neurons, with distinct morphologies, connectivity, and electrophysiological properties [1]. During embryonic mouse development, neural stem cells in the ventricular zone begin to differentiate into excitatory neurons around embryonic day 9.5 (E9.5) and adjacent ganglionic eminences to inhibitory neurons around E12.5 [2, 3]. Both kinds of neurons migrate to various brain regions, continue to establish synapses with others, and mature with their specific functions at postnatal stages. Excitatory neurons release neurotransmitters, such as glutamate, that bind to receptors on the postsynaptic neurons and cause an increase in neural activity. On the other hand, inhibitory neurons release gamma-aminobutyric acid (GABA) or glycine that decrease the likelihood that the postsynaptic neurons firing an action potential [4]. The balance between excitatory and inhibitory inputs is critical for brain function and behavior. The specification of these broad classes of neurons is largely determined by the precise regulation of gene expression networks that endow these neurons with the ability to generate different neurotransmitters, ion channels, and proteins involved in the formation of synaptic structures.

Epigenetic mechanisms including DNA methylation and histone modifications, are required to precisely regulate cell-type specific gene expression patterns. In the developing mouse brain there is a dramatic increase in genomic regions showing cell-type-specific DNA methylation [5, 6]. In neural cells, the hypomethylated genomic regions are often enriched for active histone modification markers and correlated with increased chromatin accessibility to allow the initiation and progression of RNA transcription [7]. With nuclei isolated with an affinity purification approach, highly distinctive epigenomic landscapes were reported for different types of neocortical neurons [1]. In particular, at enhancer-promoter functional domains of cell fate determining genes, chromatin structures and histone modifications are dramatically rearranged in differentiating neurons [8].

Transcription factors (TFs) are known to be essential regulators of gene expression and play critical roles in cell-fate determination. They can either bind to promoters and cooperate with transcription complex to activate transcription or bind to enhancers to elevate cell-type specific gene expression level. After neuronal induction by pro-neuronal transcription factors, the differentiation of excitatory and inhibitory neurons is controlled by a complex network of TFs in a spatiotemporal manner to achieve gene activation at specific stages during brain development. For instance, Neurogenin 2 (*Ngn2*) and Achaete-scute homolog 1 (*ASCL1*, also known as *Mash1*) are critical for the differentiation of functional excitatory neurons [9], while the LIM homeobox (*Lhx*) and distal-less (*Dlx*) family members have been shown to play a critical role in the differentiation of inhibitory neurons [10]. As neurons mature, the regulation of gene expression becomes critical for synaptic plasticity and remodeling. During brain development, transcription factors coordinate with epigenetic changes at the binding sites to ensure temporal regulation of gene expression [11]. In addition, critical transcription factors mediate changes of epigenetic signatures at their binding sites and consequently regulate gene expression. For example, EGR1, an important transcription factor in memory formation, recruits a DNA demethylase TET1 to remove the methylation marks on its binding sites and activate downstream genes [12]. Despite the growing understanding in transcription regulation at early stages of neuronal development, the contribution of transcription factor to neuro-epigenetic diversity is poorly understood.

In this study, we isolated excitatory and inhibitory neuronal nuclei from adult mouse brain to perform epigenetic comparison between these two kinds of neurons. With histone modification maps, we identified enhancer elements as the most prominent genomic regions with distinct histone modifications between excitatory and inhibitory neurons. These enhancers were predicted with abundant binding sites for a neuronal function related transcription factor, EGR1. Our additional CUT&RUN data confirmed EGR1 facilitates distinct gene regulation patterns in excitatory and inhibitory neurons.

## Results

### Generation of histone modification maps for excitatory and inhibitory neurons

To obtain nuclei from excitatory and inhibitory neurons separately, we bred Emx1-IRES-Cre knock-in (Emx1^IRES*cre*^) mice with the Sun1-tagged mice (**Figure 1A**). The Emx1^IRES*cre*^ mice express Cre recombinase in excitatory neurons and glial cells originating from the Emx1- expressing lineage, but not in GABAergic inhibitory neurons [13]. The Sun1-tagged mice contain a floxed STOP cassette, the removal of which allows the expression of nuclear membrane protein SUN1 with its C-terminus fused to a superfolder GFP (sfGFP) [1]. As such, excitatory neurons, but not inhibitory neurons, were tagged with the SUN1-sfGFP fusion protein in the resulting Sun1 f/f | Emx1-Cre (+) mice. To separate excitatory neurons from glial cells, we further stained nuclei with a fluorescent antibody targeting NeuN, a pan-neuronal marker [14]. With the combination of fluorescent signals from NeuN-immunostaining and sfGFP, we were able to remove glia cells (NeuN- and GFP+) and isolate excitatory (NeuN+ and GFP+) and inhibitory (NeuN+ and GFP-) neuronal nuclei in high purity *via* flow cytometry (**Figure 1B**). During single nuclei suspension preparation, we employed 30% iodixanol solution to remove the cell debris based on the different densities of nuclei and cell debris. This cell debris removal step allows for clean single nuclei suspension with almost no cell debris. The cell debris-free single nuclei suspension preparation was confirmed by both microscope check (**Figure S1A**) and FACS-sorting check (**Figure S1B**). The purity data was generated by FACS checking a portion of the post-sorted nuclei suspension for the same fluorescence signals (GFP and PE) (**Figure S2A-D**).

**Figure 1.**
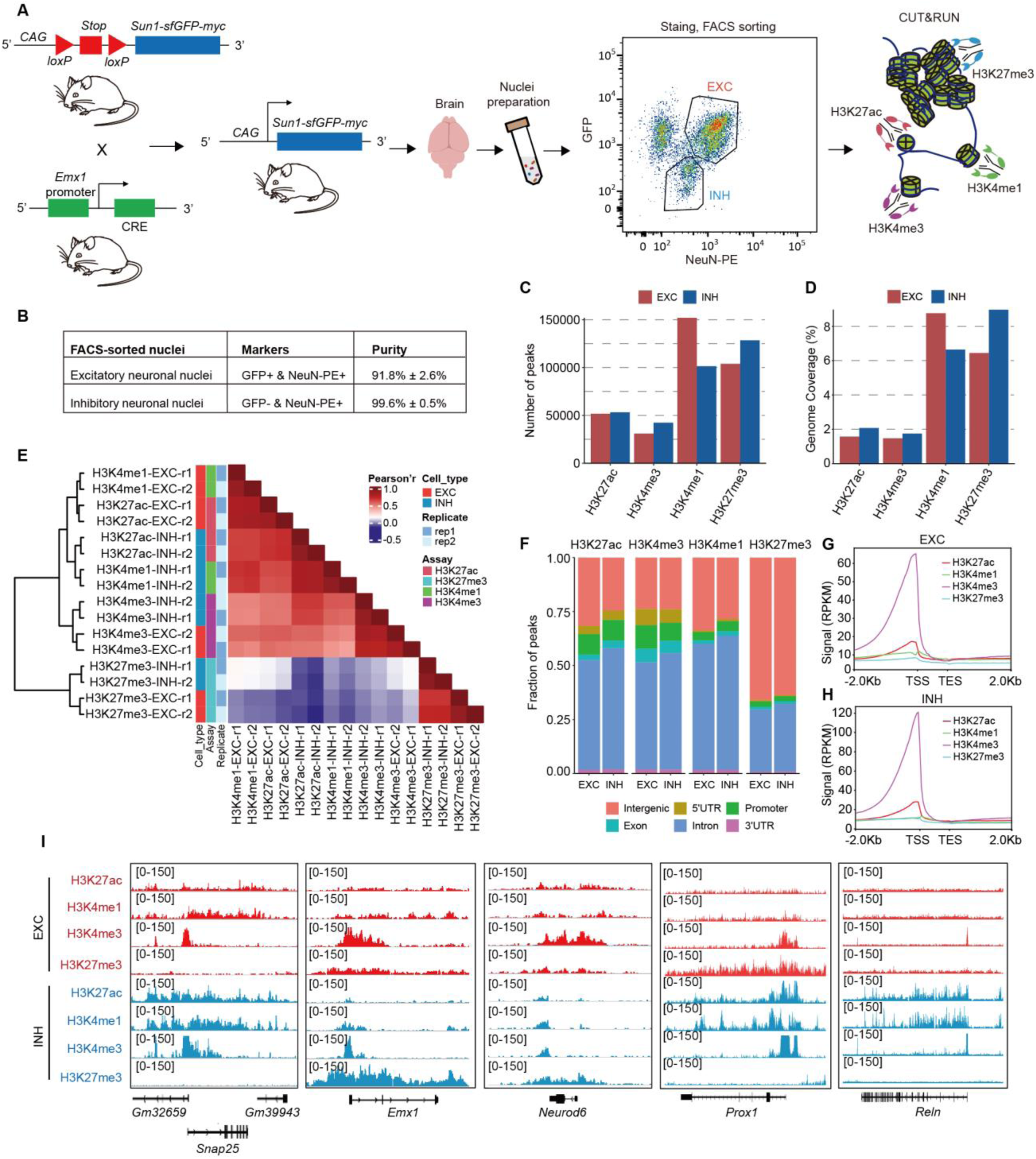
Summary of CUT&RUN datasets across neuronal subtypes and histone modifications. **A)** Diagrammatic sketch showing experimental design. **B)** Purity of the sorted nuclei for excitatory and inhibitory neurons. **C)** Numbers of reproducible peaks between biological replicates for each histone modification in excitatory and inhibitory neurons. **D)** Genome coverage of each histone modification in excitatory and inhibitory neurons. **E)** The Pearson’s correlation coefficients among histone modifications, neuronal subtypes, and biological replicates. **F)** The distribution of histone modification peaks in annotated genome. **G) and H)** Signal intensity of histone modifications surrounding TSSs in excitatory (G) and inhibitory neurons (H). **I)** Histone modification signal at well-known neuronal marker genes, including pan-neuron marker *Snap25*, excitatory neuron marker *Emx1* and *Neurod6*, inhibitory neuron marker *Prox1* and *Reln*. RPKM values in 10-bp bins are shown in each panel.

With the excitatory and inhibitory neuronal nuclei purified from adult mouse brain, we constructed CUT&RUN libraries for histone modifications including H3K27ac, H3K4me3, H3K4me1, and H3K27me3 (**Figure 1A**), with fragment sizes peaking at 168bp to 175bp (**Figure S3A**). Highly reproducible peaks (**Table S1**) between biological replicates were identified for these histone modification markers with Pearson’s R correlations in the range of 0.81 to 0.92 (**Figure S3B**). In both types of neurons, the active chromatin marker H3K27ac and the promoter marker H3K4me3 generated approximately 30,000 to 50,000 peaks, while over 100,000 peaks were identified for the enhancer marker H3K4me1 and the repressive marker H3K27me3 (**Figure 1C**). The peaks identified for these histone modifications cover around 1.6% to 9.0% of the mouse genome (**Figure 1D**). The excitatory neurons tend to have more genomic regions covered by H3K4me1 peaks while the inhibitory neurons host more repressive domains with H3K27me3 peaks. Clustering analysis of histone modification peaks confirmed the strong correlations between biological replicates (**Figure 1E**). In addition, the active chromatin marker H3K27ac and enhancer marker H3K4me1 were clustered together for both neuronal types with strong positive correlations, while negative correlations were observed between the repressive marker H3K27me3 and the rest three histone markers. We extended the clustering analysis to include a ChIP-seq dataset for excitatory neurons (**Table S2**) published in a previous study [1]. Strong correlations for H3K4me1 and H3K27ac and moderate correlations for H3K27me3 and H3K4me3 were observed for the data generated with the CUT&RUN technique in this study and the ChIP-seq procedure used previously (**Figure S4**).

We next examined the genomic distribution of the peaks for all four histone modification markers (**Figure 1F**). In general, for a given marker, peak distribution is similar between excitatory and inhibitory neurons. Not surprisingly, the active and repressive markers show striking differences in genome distribution. H3K27me3 exhibited more peaks distributed in intergenic regions, while a large fraction of H3K27ac and H3K4me3 peaks were distributed in promoter and 5’UTR regions. As expected, the promoter marker H3K4me3 showed strong intensity around transcription start sites (TSSs) followed by H3K27ac. In contrast, H3K4me1 and H3K27me3 peaks were depleted in TSSs (**Figure 1G&H**). We further scrutinized the distribution of histone modification signals for a small number of genes well-known as neuronal markers. As shown in **Figure 1I**, active histone markers were observed at the promoter site of the pan-neuron gene *Snap25* in both excitatory and inhibitory neurons. Strong signals for active markers were observed in excitatory neurons surrounding the TSSs of the excitatory neuron marker genes *Emx1* and *Neurod6*. In inhibitory neurons, strong signals for active histone markers were observed for the TSSs of *Prox1* and *Reln* genes, which are the markers of inhibitory neurons.

### Comparative histone modification analysis reveals *Egr1* as a critical transcription factor in neuronal specification

Previous studies indicated that neuronal cell specification is accompanied with substantial changes in epigenetic signatures [15, 16]. For the four histone modifications, we next determined their differential peaks between the two neuronal subtypes (**Figure S5A**). The percentages of differential peaks between excitatory and inhibitory neurons were found to be higher for the active chromatin marker H3K27ac and the enhancer marker H3K4me1, compared with the other two markers (**Figure 2A**). This result indicates that major epigenetic differences between excitatory and inhibitory neurons may occur in enhancer regions. According to the chromatin states inferred with combinations of four histone modifications, we annotated the genome into eight distinct functional regions (**Figure 2B**). The genomic regions annotated as active promoters show strong enrichment of H3K27ac and H3K4me3 and regions as active enhancers harbor more H3K27ac and H3K4me1 peaks. Not surprisingly, the histone-modification-based functional annotations were closely related to genomic annotations achieved by the distribution of known genes. For instance, active promoters annotated with histone modification were found to be overlapped with TSSs, while active enhancers, weak enhancers, and weak active domain were enriched at intergenic, intron, and 3’UTR regions (**Figure 2C**).

**Figure 2.**
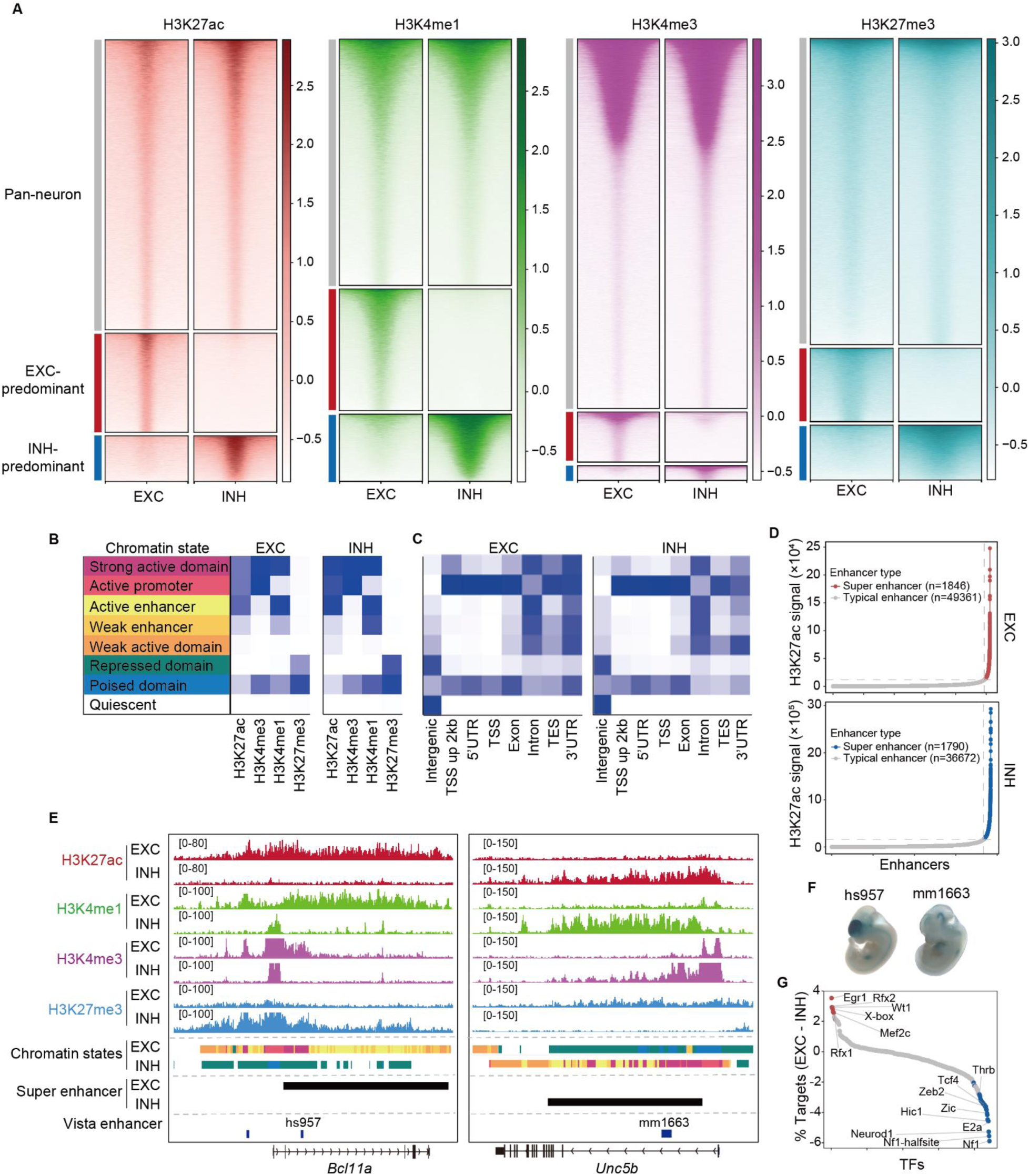
Histone modification differences between excitatory and inhibitory neurons. **A)** Heatmaps showing the difference between excitatory and inhibitory neurons for four histone modifications. Normalized z-scores were plotted in 4 kb windows centered at peaks. **B)** Eight chromatin states annotated with combinatory histone codes. **C)** Enrichment of chromatin states in annotated genome. **D)** Identification of super enhancers in neuronal subtypes. Each dot represents for an enhancer, which is classified into typical enhancer or super enhancer according to the H3K27ac signal. **E)** Examples showing histone modifications in super enhancers. **F)** Images downloaded from the VISTA database to show the reporter gene expression in transgenic mouse embryos under the control of selected enhancers. **G)** Scatterplot showing the proportion changes for TFs’ motifs at super enhancers of excitatory neurons compared with inhibitory neurons. Motifs with p value (determined by binomial test) less than 1e-65 were colored in red and blue for excitatory and inhibitory neurons, respectively.

Since super enhancers are crucial in defining cell identity [17], with these functional genome annotations, we further identified the super enhancers for the two types of neurons (**See Methods; Figure 2D & Table S3**). For example, strong H3K27ac and H3K4me1 signals were observed in the super enhancers nearby the *Bcl11a* gene in excitatory neurons but depleted in the corresponding genomic regions in inhibitory neurons (**Figure 2E**). In contrast, such a tendency was opposite in the super enhancer identified for the *Unc5b* gene. Previous studies reported that *Bcl11a* is required for neuronal morphogenesis [18] and controls the migration of cortical projection neurons [19], while *Unc5b* plays a key role in the regulation of interneuron migration to the cortex [20]. It is noteworthy that within the super enhancers identified for the *Bcl11a* and *Unc5b* genes respectively, the activities of two enhancers, hs957 and mm1663, have been validated in transgenic mouse embryos using LacZ reporters [21] (**Figure 2F**). The epigenetic states of cis-regulatory elements have an effect on the transcription factor (TF) binding and consequently regulate the expression of target genes [22, 23]. To explore the transcription factors under the influence of differential histone modifications determined in the two neuronal subtypes, we summarized and compared the TF motif frequencies in the super enhancers of excitatory and inhibitory neurons. Between the two neuronal subtypes, *Egr1*, *Rfx1*, *Rfx2* and *Mef2c* have more motifs identified in super enhancers of excitatory neurons, while *Nf1*, *Zeb2*, *Tcf4*, and *Thrb* have more potential binding sites in the super enhancers of inhibitory neurons (**Figure 2G**). Top in the ranking, *Egr1* is an immediate early response gene involving in learning and memory [24]. Our previous study showed that EGR1 binding sites are enriched in the genomic regions hypo-methylated in excitatory neurons in mouse frontal cortex [12]. Parallel analysis was performed on active promoters. *Pax7, Brn2, Chop* and *Oct11* are with motifs enriched in active promoters of excitatory neurons, while *Sp5, Boris, Klf6, Klf1* and *Znf416* have more motifs identified in active promoters of inhibitory neurons (**Figure S6**). The difference in TF motif frequencies between the promoters of two kinds of neurons is less striking when compared with that found in super enhancers.

### EGR1 favors super enhancer regions and has more binding sites detected in excitatory neurons

To explore EGR1 binding preferences in these two neuronal types, we generated EGR1 CUT&RUN libraries for excitatory and inhibitory neurons using the sorted nuclei mentioned previously (**Figure 1A&B**). A total of 24,783 and 10,391 of reproducible EGR1 peaks between biological replicates were identified in excitatory and inhibitory neurons, respectively (**Figure 3A&B; Table S4**). Interestingly, we found that EGR1 binding sites in inhibitory neurons are predominantly situated in the promoter, 5’UTR, intron, and intergenic region, while those in excitatory neurons are frequently distributed in the intron and intergenic regions (**Figure 3C**). We then annotated the EGR1 binding sites according to their chromatin states. In both types of neurons, EGR1 binding sites were depleted from the repressed domains but enriched in the active chromatin regions. In inhibitory neurons, 69.2% of EGR1 bindings sites are associated with active promoters, while this number dropped to 33.9% for excitatory neurons (**Figure 3D**). In the excitatory neurons, approximately 42.0% of EGR1 binding sites were associated with enhancers. These results were further supported by the aggregate analysis using four kinds of histone modifications, all three active histone markers were enriched at the EGR1 binding sites identified in excitatory neurons, while only H3K27ac and H3K4me3 were enriched at the EGR1 binding sites of inhibitory neurons (**Figure 3E**). To validate the result of motif analysis in previous section that EGR1 has more binding sites in the super enhancers of excitatory neurons compared with inhibitory neurons, we checked the proportion of super enhancers containing EGR1 peaks. In both neuronal types, EGR1 peaks significantly enrich at the super enhancers compared with the random genomic regions shuffled *via* 1,000 simulations (**Figure 3F; Fisher’s exact test, p value < 1e-3**). For excitatory neurons, 66.0% of super enhancers contain at least one EGR1 peak, while this number dropped to 36.3% in inhibitory neurons (Fisher’s exact test, p value < 1e-3). For instance, multiple EGR1 binding sites are located in the super enhancer surrounding *Fhl2* and *Cacng3* genes in excitatory neurons, *Prox1* and *Calb2* genes in inhibitory neurons (**Figure 3G**). Collectively, these results suggest EGR1 binding is neuronal cell type specific and in the excitatory neurons EGR1 binds more frequently to the super enhancers.

**Figure 3.**
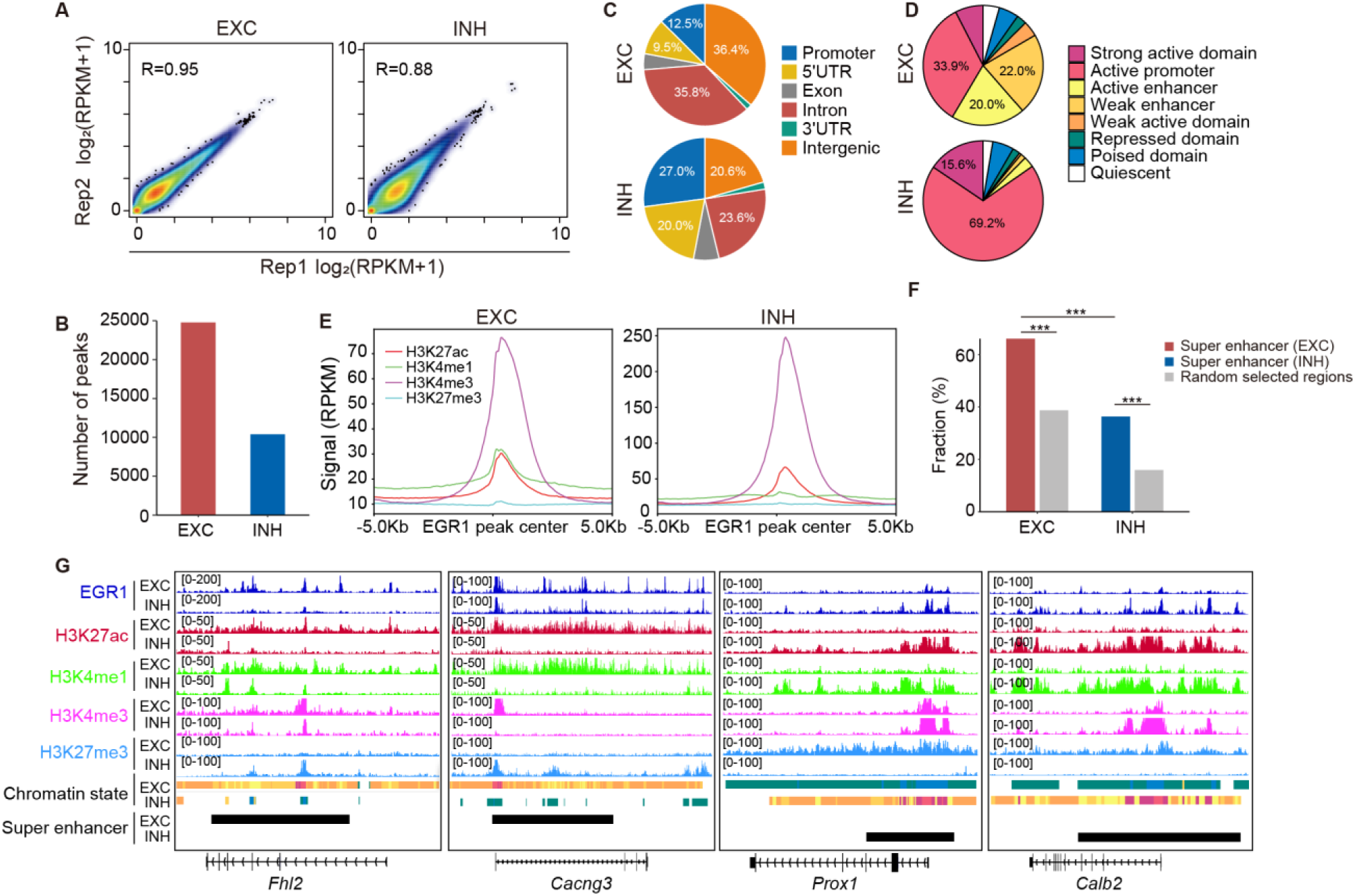
Detection of EGR1 binding sites in excitatory and inhibitory neurons. **A)** Correlations between biological replicates for EGR1 CUT&RUN datasets generated from excitatory and inhibitory nuclei, respectively. **B)** Numbers of EGR1 binding sites identified in excitatory and inhibitory neurons. **C)** Distribution of EGR1 binding sites in genome annotated with gene structure. **D)** Distribution of EGR1 binding sites across various chromatin states. **E)** Average histone modification signals around EGR1 binding sites in excitatory and inhibitory neurons. **F)** Fraction of super enhancers overlapped with EGR1 binding sites compared with those in random selected genomic regions. Statistical significance is determined by Fisher’s exact test, the comparisons with p value less than 1e-3 are labeled with ***. **G)** Examples showing EGR1 binding sites and histone modifications around super enhancers.

To further investigate EGR1 binding difference in two neuronal subtypes, we set the two-fold changes in peak signals as the threshold of differential binding (**Figure S5B**). Among the 26,686 EGR1 peaks in the two types of neurons, 6,506 and 1,907 peaks were identified as EXC- predominant and INH-predominant respectively (**Figure 4A**). The rest of the 18,273 peaks were annotated as pan-neuronal EGR1 binding sites in both types of neurons. These 18,273 EGR1 binding sites share similar epigenetic features between excitatory and inhibitory neurons, including chromatin accessibility, DNA methylation profile (**Table S2**), and histone modifications. In particular, in both neuronal types, the three active histone markers H3K27ac, H4K4me1 and H3K4me3 were observed in pan-neuronal EGR1 binding sites. In contrast, for the EXC-predominant EGR1 peaks, stronger signals of H3K27ac and H3K4me1 were observed in excitatory neurons. Interestingly, for INH-predominant EGR1 peaks, stronger H3K27ac and H3K4me3 signals were observed in inhibitory neurons (**Figure 4A**). These results suggest the EXC- predominant and INH-predominant EGR1 peaks may have different roles in these distinct neuronal populations. EXC-predominant EGR1 peaks may serve as enhancers while the INH-predominant peaks may have promoter activity. Although slightly weaker than those in inhibitory neurons, H3K4me3 signal in excitatory neurons is observed surrounding INH-predominant peaks.

**Figure 4.**
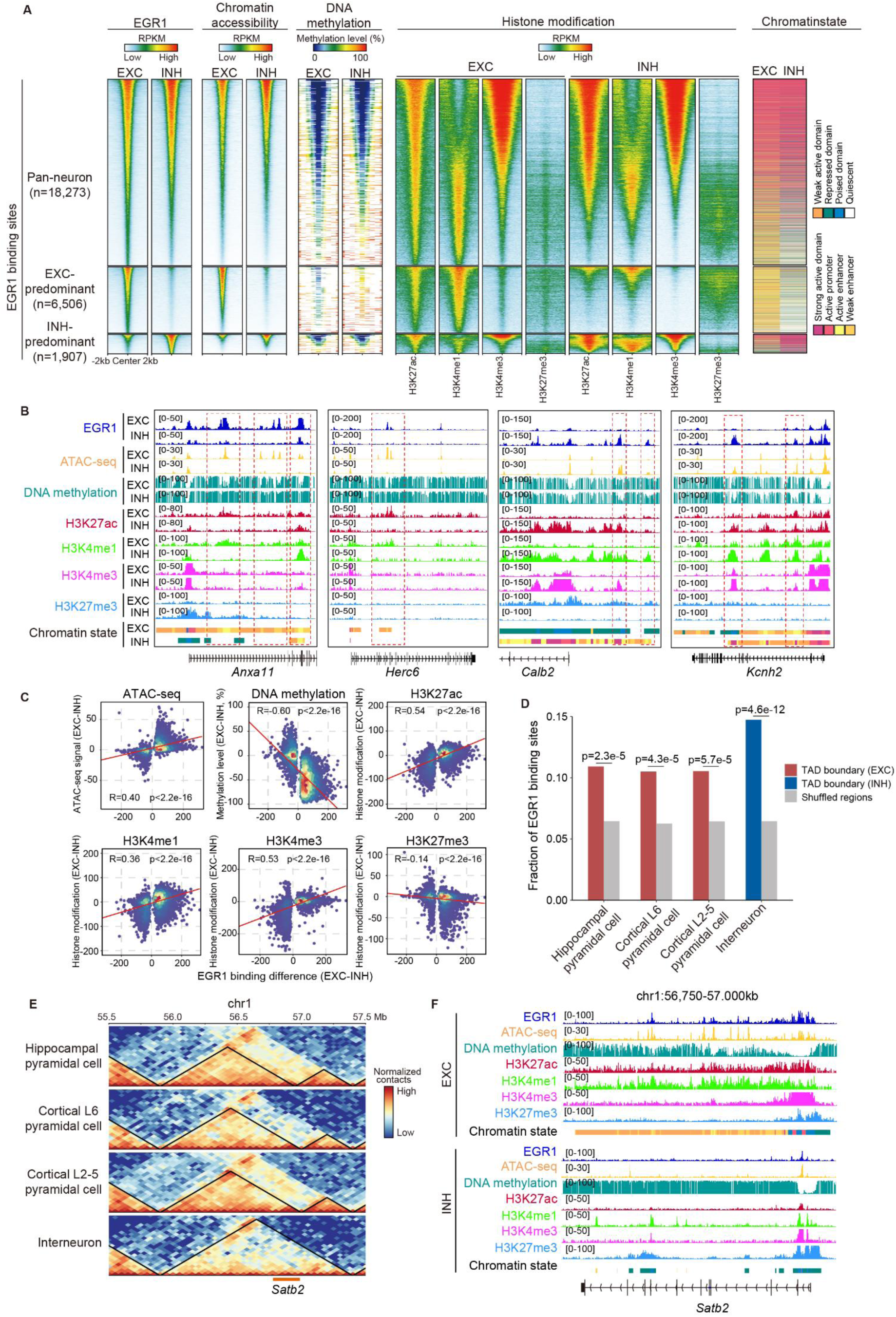
Epigenetic signatures of EGR1 binding sites. **A)** Epigenetic marks plotted in 4 kb windows centered at EGR1 binding sites. **B)** Selected examples showing neuronal subtype- predominant EGR1 binding, chromatin states and epigenetic signatures. **C)** Correlations between EGR1 binding and various epigenetic markers. **D)** Fraction of EGR1 binding sites located in the TAD boundary of various cell types compared with randomly selected regions via 1000 times shuffle. Statistical significance is determined by Fisher’s exact test, the p values are labeled at the top of each comparison. **E)** Contact maps of multiple neuronal subtypes at 50-kb resolution showing EGR1 binding sites in excitatory neuron-specific TAD boundary at *Satb2* gene locus. **F)** EGR1 binding signal and histone modifications at *Satb2* genes adjacent to an excitatory neuron specific TAD boundary showing in (E).

To further explore the association between EGR1 binding and other epigenetic markers, we re-analyzed ATAC-seq and MethylC-seq data for excitatory and inhibitory neurons generated in a previous study [1] (**Table S2**). Strong EGR1 peaks are accompanied with high chromatin accessibility and low DNA methylation level (**Figure 4A&B**). This result suggests that EGR1 tends to bind to active chromatin regions. We then calculated the Pearson’s correlation coefficients between EGR1 binding and various epigenetic markers (**Figure 4C**). The correlation between EGR1 binding and DNA methylation is -0.60. This is consistent with the fact that EGR1 is able to recruit DNA demethylation enzyme TET1 to its binding sites [12]. As expected, strong correlations were observed between EGR1 binding and multiple epigenetic markers for open chromatin regions. Among all four histone modifications, H3K27ac shows the strongest correlation (Pearson’s r = 0.54) while H3K27me3 shows a negative correlation (Pearson’s r = -0.14) with EGR1 binding.

Numerous data demonstrate that the three-dimensional (3D) genome structure plays an important role controlling the interaction of genomic DNA with transcription factors to achieve gene expression regulation [8, 25, 26]. A previous study reported that *Egr1* motifs are more abundant at pyramidal glutamatergic neuron-specific contacts compared with dopaminergic neurons [27]. To check whether EGR1 binding is associated with 3D chromatin conformation, we re-analyzed the aggregated single-cell diploid chromatin conformation capture (Dip-C) data from three excitatory neurons (cortical layer 2-5 pyramidal cells, cortical layer6 pyramidal cells and hippocampal pyramidal cells) and interneurons [28] (**Table S2**), and inferred the topologically associated domain (TAD) of these neurons. We found 10.5% and 14.7% of EGR1 binding sites in excitatory and inhibitory neurons are located in their TADs respectively, which are significantly higher than the fraction of those in randomly selected regions (**Figure 4D**), indicating that EGR1 may contributes to the genome architecture formation in neuronal subtypes. For instance, at the *Satb2* gene locus (which is a determinant for upper-layer neuron specification) and its upstream region, EXC-predominant EGR1 binding sites were observed at the excitatory-specific TAD boundary associated with multiple active epigenetic markers (**Figure 4E&F**). Collectively, our results demonstrated that EGR1 binding is highly correlated with active epigenetic markers and may contribute to cell-type specific chromatin architectures.

### Differential EGR1 binding is associated with distinct gene pathways and expression program in the two neuronal subtypes

To explore the functional relevance of differential EGR1 binding in excitatory and inhibitory neurons, we inferred the EGR1 target genes by utilizing genomic annotation and 3D chromatin structure. The genes with EGR1 peaks in the promoter, gene body, and simultaneously within the same topologically associated domain (TAD) were defined as EGR1 target genes (**Figure 5A**). With EXC- and INH-predominant EGR1 peaks, 3,183 and 1,107 genes were identified as EGR1 target genes, respectively. With neuronal cell-type specific RNA-seq datasets [1], we found that *Egr1* was strongly expressed in excitatory neurons (fold change=1.53) compared to inhibitory neurons (**Figure 5B**). Since EGR1 may recruit TET1 to remove the methylation marks and activate downstream genes [12], it may serve as a positive regulator of its target genes. As expected, genes with EXC-predominant EGR1 peaks showed slightly but significantly higher expression levels in excitatory neurons compared to those in inhibitory neurons, and *vice versa* (**Figure 5C**). For example, *Cacng3, Fhl2, Herc6, Anxa11* and *Satb2* genes associated with EXC-predominant EGR1 binding sites were upregulated in excitatory neurons, while INH-predominant EGR1 binding sites associated genes such as *Prox1, Calb2* and *Kcnh2* were upregulated in inhibitory neurons (**Figure 5D**). To further explore the regulatory functions of EGR1 in excitatory and inhibitory neurons, we performed the KEGG pathway enrichment analysis for genes associated with EXC- and INH- predominant EGR1 binding sites. Genes with EXC-predominant EGR1 peaks are enriched in “Axon guidance”, “Calcium signaling pathway” and “Glutamatergic synapse”. Genes with INH- predominant EGR1 peaks are enriched in “Dopaminergic synapse”, “Neurotrophin signaling pathway” and some disease pathways, such as “Neurodegeneration-multiple diseases”, “Huntington disease” and “Alzheimer disease” (**Figure 5E**). In summary, these results reveal that EGR1 is involved in neuronal specification and plays distinct roles in neuronal subtypes.

**Figure 5.**
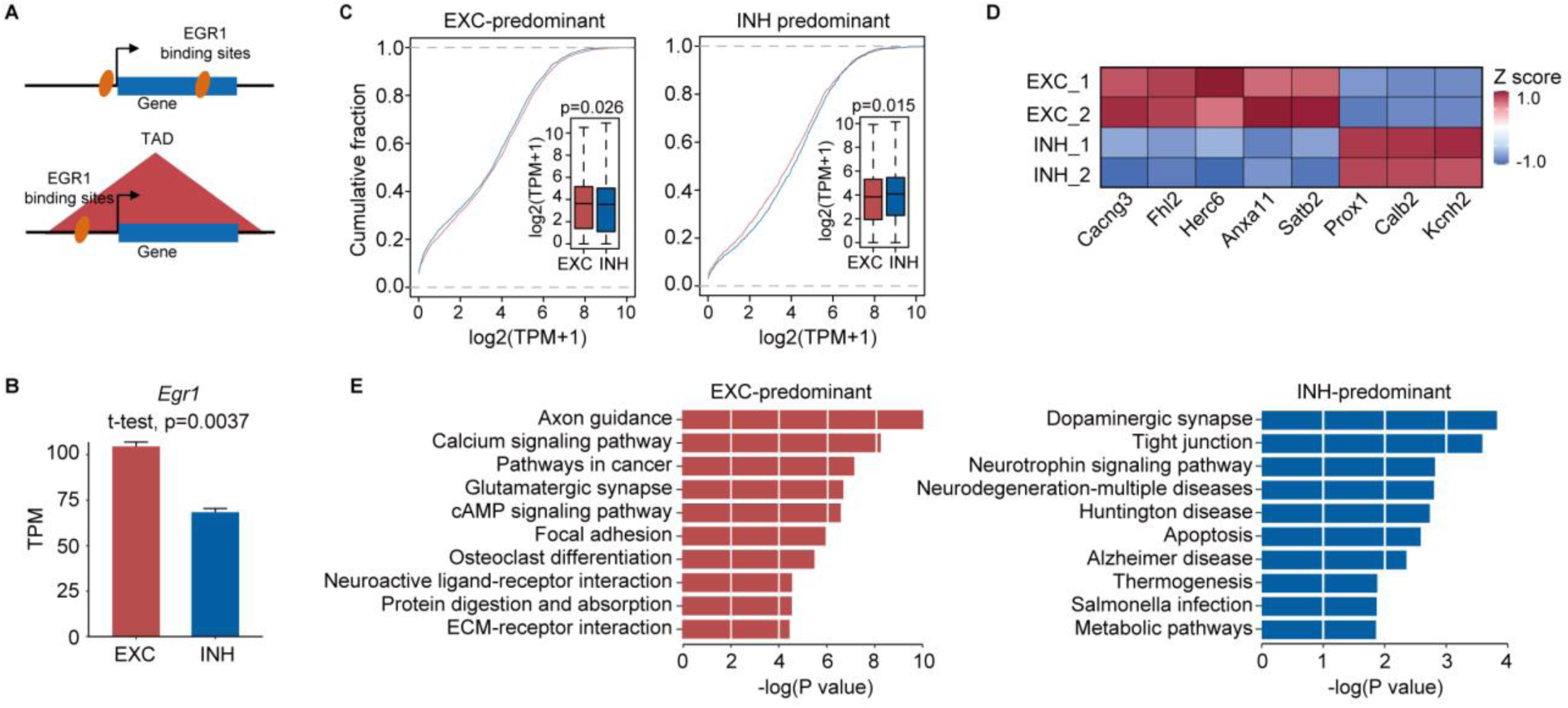
Functional characterization of neuronal-subtype predominant EGR1 binding sites. **A)** Illustration of EGR1 target gene inference. **B)** Expression of *Egr1* in excitatory and inhibitory neurons. **C)** Expression cumulative curve of neuronal subtype-predominant EGR1 binding sites associated genes. Statistical significance was determined by Mann-Whitney U test. **D)** Expression of selected examples in excitatory and inhibitory neurons. **E)** KEGG pathway analysis for neuronal subtype-predominant EGR1 binding sites associated genes.

### Establishment of EGR1 regulatory networks differs in two neuronal subtypes during brain development

Since EGR1 binding is strongly correlated with active epigenetic markers (**Figure 4C**), the changes in chromatin accessibility of EGR1 binding sites could reflect dynamic EGR1 binding. To illustrate how the epigenetic landscape of EGR1 binding sites was established during brain development, we made use of single-nucleus ATAC-seq (snATAC-seq) datasets generated with developing mouse brains from E12.5 to P56 [29, 30]. We focused on excitatory and inhibitory populations of neurons for these analyses (**Figure 6A**). Aggregated ATAC-seq data demonstrated the opening chromatin states of a pan-neuronal gene *Snap25* in both neuronal types, while neuronal cell-type-specific genes such as *Neurod6* and *Dlx5* were only accessible in excitatory and inhibitory neurons, respectively (**Figure S7A**), which confirmed that these aggregated ATAC-seq data could effectively reflect the neuronal specificity. We next calculated the number of accessible EGR1 peaks in each stage. Although this number increased in both neuronal types during brain development, considerable fraction of INH-predominant EGR1 peaks was activated at early stages, while EXC-predominant EGR1 peaks gain accessibility gradually during brain development and such a trend accelerates in postnatal stages (**Figure 6B**). According to the aggregated snATAC-seq signal of each stage, the majority of pan-neuron and almost all INH-predominant EGR1 peaks become accessible before P0, while most EXC-predominant EGR1 peaks gain strong signal in postnatal stages (**Figure 6C**). For example, EXC-predominant EGR1 peaks surrounding *Anxa11, Herc6, Fhl2* and *Cacng3* become accessible only after P0 in excitatory neurons, while INH- predominant EGR1 peaks surrounding *Calb2* and *Kcnh2* become accessible in early embryonic stages in inhibitory neurons (**Figure 6D&E**). These results suggested that the accessibility of neuronal cell-type-specific EGR1 peaks may be established at different time points during brain development.

**Figure 6.**
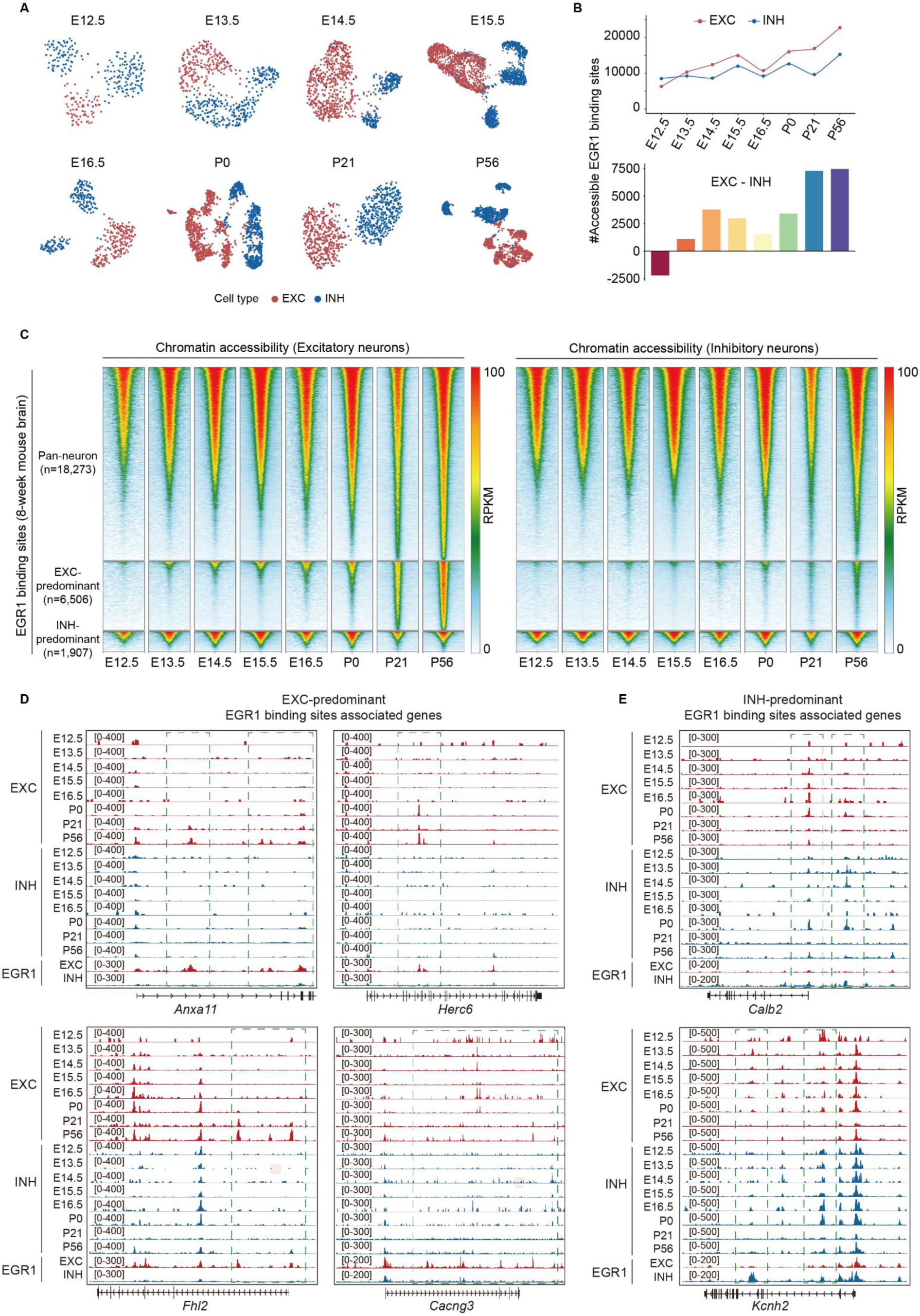
Chromatin accessibility of EGR1 binding sites in excitatory and inhibitory neurons during brain development. **A)** Single-nucleus ATAC-seq (snATAC-seq) data analysis to show the umaps of excitatory and inhibitory nuclei during brain development. **B)** Number of accessible EGR1 binding sites during brain development. Top panel showed the number of accessible EGR1 binding sites in excitatory and inhibitory neurons, respectively. Bottom panel showed the number difference of accessible EGR1 binding sites between excitatory and inhibitory neurons. **C)** Heatmap showing chromatin accessibility of EGR1 binding sites during brain developments. **D and E)** Examples to show the chromatin accessibility of EXC- (D) and INH-predominant (E) EGR1 binding sites during brain development.

To further understand the establishment of EGR1 regulatory network, we explored the expression of *Egr1* together with its target genes in excitatory and inhibitory neurons during brain development using single-cell RNA-seq (scRNA-seq) datasets generated from E12.5 to P60 mouse brains [31–33]. Gene expression data for excitatory and inhibitory neurons were extracted for downstream analysis (**Figure 7A&B**). Integrative analysis was preformed to merge scRNA-seq with snATAC-seq datasets for each development stage. We observed that the neuronal subtypes identified using two kinds of datasets were comparable (**Figure S7B**). To explore the dynamic gene expression for the two neuronal types during brain development, aggregated analysis of scRNA-seq data generated for each stage was performed. For example, *Snap25* was expressed in both neuronal populations and dramatically increased in postnatal stages, while *Neurod6* and *Dlx5* were expressed in excitatory and inhibitory neurons, respectively (**Figure S7C**). To examine whether these datasets can be successfully integrated, we identified the top 200 specifically expressed genes in each stage for two neuronal types and checked their functions, gene ontology results showed that development related terms, such as “axonogenesis”, “synapse organization” and “axon guidance” were enriched in embryonic stages, and neuronal related functions, such as “neurotransmitter transport” and “synaptic vesicle cycle” were enriched in postnatal stages (**Figure S7D**). Despite these “omics” data were generated by different labs, the successful data integration enables us to provide a continuous view of brain gene expression and chromatin accessibility from embryonic stage E12.5 to postnatal P56.

**Figure 7.**
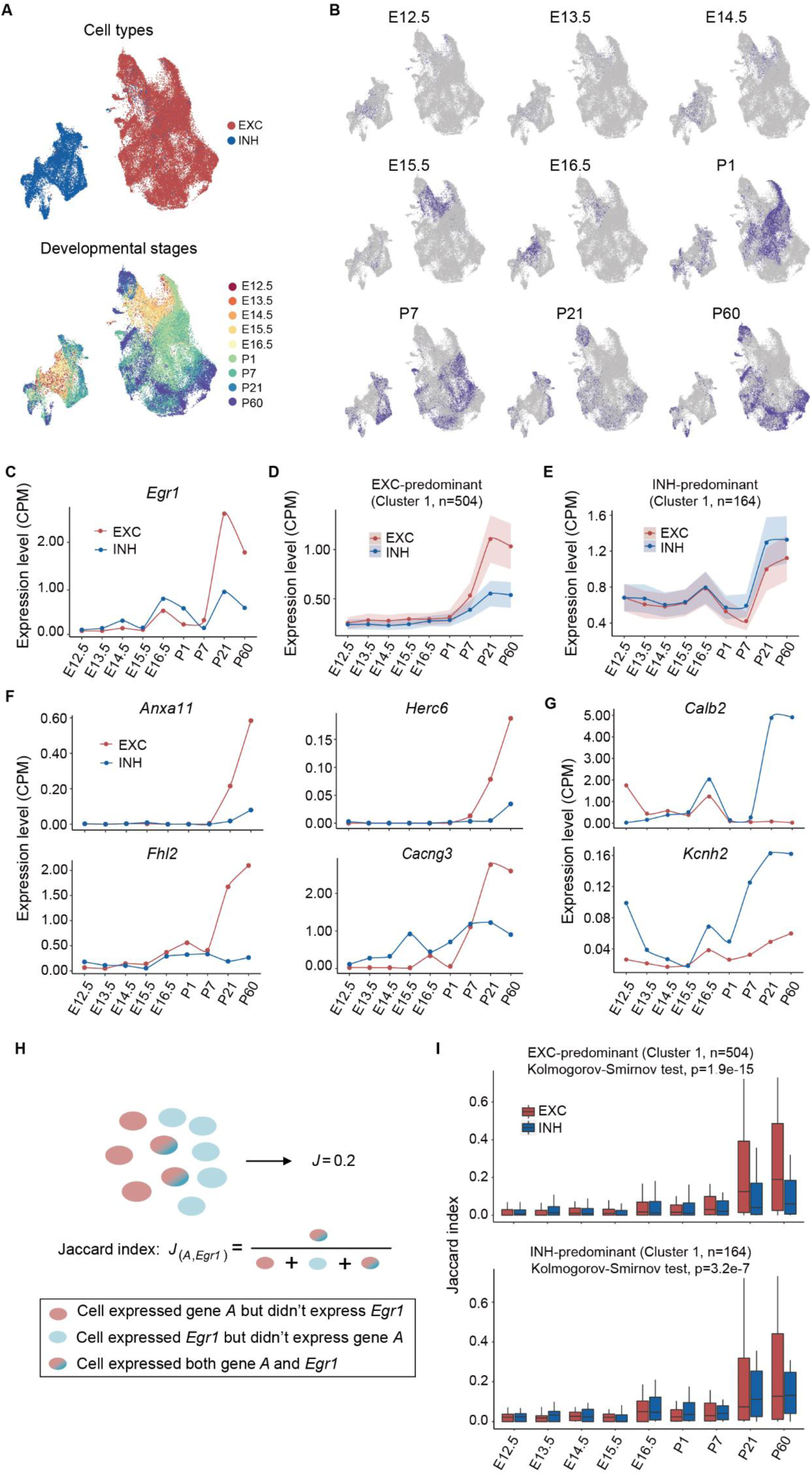
Single-cell RNAseq analysis of EGR1 target genes during mouse brain development. **A)** Umaps showing the excitatory and inhibitory neurons across mouse brain development. **B)** Umaps showing the neurons in each developmental stage. **C)** *Egr1* expression in excitatory and inhibitory neurons during brain development. **D)** Average expression of cluster 1 of EXC- predominant EGR1 peaks associated genes during brain development. **E)** Average expression of cluster 1 of INH-predominant EGR1 peaks associated genes during brain development. **F and G)** Examples to show the expression of genes associated with EXC- (F) and INH-predominant (G) EGR1 peaks during brain development. **H)** Sketch to show the co-expression between *Egr1* and its target genes. **I)** Jaccard index between *Egr1* and its target genes during brain development in excitatory and inhibitory neurons.

The expression of the *Egr1* gene increases during brain development in both neuronal types, especially in postnatal stages, the expression level of *Egr1* in excitatory neurons was over two-fold higher than inhibitory neurons at P21 and P60 (**Figure 7C**). To examine the effect of EGR1 on its target genes during brain development, we performed the clustering analysis for genes associated with EXC- and INH-predominant EGR1 binding sites respectively (**Figure S8A&B**). A large number of EGR1 target genes show various kinds of expression patterns distinct from that of *Egr1*, which suggests other mechanisms may participate in the regulation of these genes. Interestingly, 504 and 164 genes among EXC- and INH-predominant EGR1 peaks associated genes respectively were found to share similar expression pattern with *Egr1* (**Figure 7D&E**). For example, the expression of *Anxa11, Herc6, Fhl2, Cncng3* in excitatory neurons, *Calb2* and *Kcnh2* in inhibitory neurons were synchronized with *Egr1* during brain development (**Figure 7F&G**). The 504 genes share similar expression pattern with *Egr1* enriched in “Calcium signaling pathway”, “Cholinergic synapse” and “Glutamatergic synapse” in excitatory neurons. In inhibitory neurons, the 164 genes share similar expression pattern with *Egr1* enriched in “Metabolic pathways” and “Huntington disease” (**Figure S8C&D**). Additionally, we examined the co-expression relationship between these genes and *Egr1* across neuronal types during brain development using the Jaccard index (**Figure 7H**). While the Jaccard index increased along with brain development in both neuronal subtypes, genes associated with EXC-predominant EGR1 peaks in excitatory neurons exhibited higher expression levels than those in inhibitory neurons, particularly in postnatal stages, and vice versa (**Figure 7I**). Overall, these findings suggest that *Egr1* may play a critical role in regulating the expression of these gene subsets, serving as a key regulator in neuronal specification.

## Discussion

Current understanding of brain epigenetic regulatory network is still very limited, in particular for the link between epigenetic programming and neuronal specification. Only a handful of datasets have been generated to demonstrate the roles of histone modification and DNA methylation in controlling chromatin loops mediated by transcription factors in a cell type specific manner. In a previous study, Mo et al generated a comprehensive epigenome dataset for excitatory and inhibitory neurons, including methylomes, ATAC-seq data, and ChIP-seq data for histone modifications [1]. Their comparative analysis provided a link between epigenomic diversity with the functional and transcriptional complexity of neurons. Due to the low proportion of inhibitory neurons, only histone modification maps for excitatory neurons were obtained at that time. Recently, single-cell epigenetics technologies help in gaining insight into the cell-type specific gene regulatory programs. Zhu et al developed a Paired-Tag method for joint profiling of histone modifications and transcriptome in single cells, and applied it to frontal cortex and hippocampus of adult mice to produce cell-type-resolved maps of chromatin state and transcriptome [16]. Despites these advances, it remains largely unknown that how cell-type specific gene regulation is achieved for neuron subtypes.

In this study, we provided genome-wide chromatin state maps with high-coverage CUT&RUN data of four kinds of histone modification for excitatory and inhibitory neurons. Our CUT&RUN data of histone modifications in excitatory neurons is highly comparable with the ChIP-seq data generated by Mo et al. The comparative histone modification analysis demonstrated that, between two kinds of neurons, the percentages of differential peaks identified for H3K4me1 and H3K27ac are higher than those for H3K4me3 and H3K27me3. This indicates that cell-type specific histone modification is primarily enriched in enhancers. Despite that some studies have demonstrated the functional importance of enhancers in brain cell types and examined its relationship with disease-risk [34, 35], our further motif analysis in super enhancer regions of excitatory and inhibitory neurons identified EGR1 as critical transcription factor involves in neuronal specification.

Our previous study indicated that EGR1 is able to recruit DNA demethylation enzyme TET1 to activate downstream gene expression during postnatal brain development [12]. In addition, DNA demethylation mediated by EGR1 is largely limited to excitatory neurons but how EGR1 achieves cell-subtype specific functions remains unknown. In this study, our EGR1 CUT&RUN data indicated that EGR1 mediates distinct programs of gene expression regulation in two kinds of neuron and involves in the formation of super enhancers in excitatory neurons. The neuronal cell-subtype specific EGR1 binding is associated with distinct epigenetic signatures including DNA methylation, chromatin accessibility, histone modifications, and 3D genomic conformation. Such interplay between epigenetic marks and transcription factors may be generalized to other neurodevelopmental processes or other cell types. Although the cause-effect relationship between histone modifications and EGR1 binding remain unexplored, this study provided a comprehensive set of histone maps together with EGR1 binding profiles and shed lights on distinct epigenetic regulation between excitatory and inhibitory neurons.

## Conclusion

Our comprehensive histone modification and CUT&RUN data in neuronal types supported that EGR1 serves as a key regulator in neuronal specification through epigenetic mechanisms. These findings provide valuable insights into the mechanisms underlying neuronal subtype specification and establish a framework for future investigations into cell type-specific transcriptional regulation in the nervous system.

## Methods

### Mice

The animal experiments had been approved prior to the study by the Institutional Animal Care and Use Committee (IACUC) of Virginia Tech. Mice were maintained and bred in a 12-hour light/dark cycle under standard pathogen-free conditions. The Sun1 mice (strain #: 021039) and Emx1-IRES-Cre mice (strain #: 005628) were obtained from Jackson laboratory. Crude DNA was extracted from tail biopsies using Direct PCR tail lysis buffer supplemented with Proteinase K solution and genotyped by PCR according to the Jackson Laboratory’s protocols. Adult (8 weeks) male mouse brain samples were used for experiments.

### Nuclei isolation

Mice were euthanized by inhalation of carbon dioxide (CO2). Cervical dissociation was further performed and the brain tissues were rapidly dissected. Nuclei preparation was adapted from previous publications [36, 37]. Briefly, the mouse brain tissue was Dounce homogenized in NE buffer (0.32M sucrose, 10 mM Tris-HCl pH 8.0, 5 mM CaCl2, 3 mM MgCl2, 1 mM DTT, 0.1 mM EDTA, 0.1% Triton X-100, 1x Proteinase Inhibitor Cocktail), incubated on ice for 10min, filtered through 70 μm cell strainer (Miltenyi Biotec, cat# 130-098-462), and spun down at 1000g for 5min at 4°C. The supernatant was removed and the pellet nuclei was further purified using a 30% iodixanol cushion and centrifuged at 8,000g for 20min at 4°C. The cell debris on the top of the supernatant were aspirated, and the purified nuclei were pelleted at the bottom of the tube.

### Nuclei staining and FACS sorting

The purified nuclei were resuspended in PB buffer (1xPBS with 1% BSA, 1x Proteinase Inhibitor Cocktail) and incubated with mouse anti-NeuN-PE antibody (Sigma, cat# FCMAB317PE) for 1h at 4°C. The stained nuclei were washed twice with PB buffer, resuspended in PB buffer, and subjected to FACS sorting procedures using the BD FACS ARIA Flow Cytometer. Both excitatory neuronal nuclei (GFP+ and NeuN+) and inhibitory neuronal nuclei (GFP- and NeuN+) were collected.

### Cleavage Under Targets and Release Using Nuclease (CUT&RUN)

CUT&RUN was performed using the CUTANA CUT&RUN Kit (EpiCypher, Cat# 14-1048) as previously described [38]. Briefly, the ConA Beads were washed twice and resuspended in Bead Activation Buffer. The FACS-sorted nuclei were pelleted and resuspended in Wash Buffer and mixed with the ConA Beads. The nuclei-bead slurry was incubated on a tube rotator for 10min at room temperature (RT), allowing the nuclei absorbed to the beads. The nuclei/beads conjugates were resuspended in 50 μL of Antibody Buffer (Wash Buffer with 0.01% Digitonin and 2 mM EDTA) containing 2 μg of H3K27ac antibody (abcam, cat# ab4729), or H3K4me1 antibody (Active Motif, cat# 39498), or H3K4me3 antibody (Active Motif, cat# 39060), or H3K27me3 antibody (Active Motif, cat# 39055), or EGR1 antibody (Santa Cruz, cat# sc101033) and incubated in a tube nutator overnight at 4°C. The next morning, the nuclei/beads conjugates were washed twice in 200 μL of Cell Permeabilization Buffer (Wash Buffer with 0.01% Digitonin), resuspended in 50 μL of Cell Permeabilization Buffer, and 2.5 μL pAG-MNase (20x stock) was added. The nuclei/beads conjugates were incubated for 10min at RT, followed by two washes in 200 μL of Cell Permeabilization Buffer, and resuspension in 50 μL of Cell Permeabilization Buffer. Tubes were chilled on ice, 1 μL of 100 mM Calcium Chloride were added, and the tubes were nutated for 2h at 4°C. Then 33 μL of Stop Buffer and 1 μL of Spike-in DNA (0.5ng/μL) were added to each tube. The tubes were incubated for 10min at 37°C, and placed on a magnet stand until slurry cleared. The supernatant containing CUT&RUN enriched DNA fragments were collected in 1.5mL tubes and DNA purification was performed using the DNA Cleanup Columns provided in the kit following the manufacturer’s instructions.

### Construction and Sequencing of CUT&RUN Libraries

Libraries for CUT&RUN samples were prepared using the NEBNext Ultra II DNA Library Prep Kit for Illumina (NEB, cat# E7645S) following the manufacturer’s instructions. Briefly, the CUT&RUN enriched DNA fragments were end-repaired and dA-tailed, and ligated to DNA adaptors. After purification with Ampure beads, PCR amplification was performed to enrich adaptor-ligated DNA fragments. Molar concentration of the finished libraries was estimated using a combination of Qubit dsDNA HS assay kit (Thermo Fisher, cat# Q32854) on Qubit 3.0 Fluorometer (Thermo Fisher, cat# Q33218) and Agilent DNA D1000 Screen Tape (Agilent, cat# 5067-5582) on 4150 TapeStation System (Agilent, cat# G2992AA). Individually indexed libraries were pooled and sequenced on Novaseq 6000 platform with paired end 150bp mode.

### CUT&RUN data analysis

For all reads derived from cun&run libraries, sequencing adapters and low-quality bases were first trimmed with cutadapt (v1.18, https://github.com/marcelm/cutadapt/) and trim_galore (v0.5.0, https://www.bioinformatics.babraham.ac.uk/projects/trim_galore/). The retained reads were aligned to mouse genome (mm10) using bowtie2 (v2.3.5) [39] in pair-end mode with option “-N 1 -L 25”. PCR duplications were removed using picard with the option “REMOVE_DUPLICATES=true” (v2.25.0, https://broadinstitute.github.io/picard/). Non- redundant reads were further filtered for minimal mapping quality (MAPQ ≥ 30) using samtools view (v1.12) [40] with option “-q30”.

Peak calling for histone modifications was performed by MACS2 (v2.2.5) [41] using option “-p 0.05” for H3K27ac and H3K4me3 and “--broad -p 0.05” for H3K4me1 and H3K27me3. The reproducible peaks between biological replicates were further identified following irreproducible discovery rate (IDR, v2.0.4.2) framework [42] with parameters “--rank signal.value”. Stricter parameters were adopted for EGR1 cun&run datasets peak calling to generate the highly reliable transcription factor binding sites, with option “-p 0.005” for MACS2 and “--rank signal.value -- idr-threshold 0.02” for IDR framework.

### The clustering analysis and correlation among neuronal subtypes, markers and biological replicates

The correlation coefficient between samples was calculated as following: the RPKM value was generated on a 1Kb-window base, the signal score was then summed within each 5kb-window for the entire genome and was compared across different samples. Pearson correlation coefficient was used for all analyses and hierarchical clustering was adopted for clustering analysis.

### Detection of neuronal-subtype predominant histone modification peaks and EGR1 binding sites

DESeq2 (v1.30.1) [43] was adopted to perform differential peak analysis between excitatory and inhibitory neurons for H3K27ac, H3K4me3, H3K4me1, H3K27me3 and EGR1, separately. For each marker, firstly, a union peak list of excitatory and inhibitory neurons was generated by bedtools merge (v2.30.0) [44]. The read count in the union peak regions were then calculated and normalized by “count” function in the DESeq2. Finally, the differential peak regions were determined by “result” function in the DESeq2, with thresholds “padj≤0.05” and “FoldChange≥ 2” or “FoldChange≤0.5”.

### Annotation of chromatin states

ChromHMM (v1.23) [45] was adopted to annotate the chromatin states. In brief, BinarizeBam function was first used to divide the mouse genome into 200bp non-overlapped bins and convert the signal in bam file to binary data in 200bp bins for each histone modification marker in two neuronal subtypes, respectively. The biological replicates were merged and considered as one sample. LearnModel function was then used to train the prediction model by integrating the four histone modification markers and assign the 200bp bins into multiple chromatin states. The 8-state model was selected, since it presented maximum number of chromatin states with distinct histone modification marker combinations. The 8 chromatin states were labeled based on their combinations of histone modifications.

### Identification of super enhancers

Ranking of super enhancer (ROSE) [17] was used to identify super enhancers. Genomic regions annotated as “active enhancer”, “weak enhancer”, and “strong active domain” were selected and merged as an enhancer pool. The enhancers in the pool located within 12.5 kb from each other were merged and then ranked by the H3K27ac signal. The point with the tangent slope equals to 1 was selected as the inflection point to classify super enhancers and typical enhancers. Enhancers above the point were defined as super enhancers and the rest were defined as typical enhancers.

### Motif analysis

Homer software [46] was applied to perform motif analysis. “findMotifsGenome.pl” function was used to search all the motifs in each genomic sequence for super enhancers of excitatory and inhibitory neurons, respectively. For each motif, the percentage of enhancers containing this motif was calculated for excitatory and inhibitory neurons, separately. The binomial test was used to determine the statistical significance of the percentage difference in two neuronal subtypes for each motif.

### Re-analysis of RNA-seq, MethylC-seq, ATAC-seq, ChIP-seq and Dip-C data

MethylC-seq and ATAC-seq data of three sorted neuronal subtypes from adult mouse neocortex were downloaded from previous study [1], including excitatory (EXC) neurons, parvalbumin (PV) expressing fast-spiking interneurons, and vasoactive intestinal peptide (VIP) expressing interneurons (**Table S2**), each sample with two biological replicates. The data of PV and VIP neurons were merged as inhibitory neurons.

For MethylC-seq datasets, sequence adapters and low-quality bases were filtered with cutadapt and trim_galore. The retained reads were aligned to mouse genome (mm10) using bismark [47] with default parameters, PCR duplications were removed using deduplicate_bismark module embedded in bismark software, and genome-wide cytosine methylation report was generated by using bismark_methylation_extractor module in bismark software. The CpG dinucleotides covered by at least 10 reads were retained for downstream analysis.

For RNA-seq datasets of excitatory and inhibitory neurons, adapters and bases of low quality were trimmed and the remaining reads were mapped to the mouse genome (mm10) by RSEM [48] with Bowtie2 to achieve the expression level of each gene. TPM (Transcripts Per Million) values were adopted for downstream analysis.

For ATAC-seq datasets, quality control was performed using the same strategy with MethylC- seq datasets. The retained high-quality sequences were aligned to mouse genome (mm10) using bowtie2 with parameter “-N 1 -L 25”. The average RPKM value in non-overlapped 10-bp bins were calculated and used for downstream analysis.

ChIP-seq of H3K27ac, H3K4me1, H3K4me3 and H3K27me3 for sorted excitatory neurons from adult mouse brain were downloaded from previous study [1]. Quality control was performed using the same pipeline with MethylC-seq and ATAC-seq datasets. The retained high-quality sequences were aligned to mouse genome (mm10) using bowtie2 with parameter “-N 1 -L 25”.

Aggregated scDip-C contact matrix for cortical layer 2-5 pyramidal cells and interneurons were downloaded from GEO datasets with accession GSE146397. HiCExplorer [49] was adpoted for data analysis, hicNormalize function was used to normalize the contact matrix, hicFindTADs function was used to detect topologically associated domain (TAD).

### Functional enrichment analysis

KEGG pathway enrichment analysis was performed by DAVID web server [50] with default parameters. The gene symbols of EGR1 target genes were input to server, fisher’s exact test was used to perform enrichment analysis.

### Analysis of snATAC-seq data

snATAC-seq data from developing mouse brain at E12.5, E13.5, E14.5, E15.5, E16.5, P0, P21 and P56 were downloaded [29, 30] (**Table S2**) and re-analyzed following the instructions in previous study [29] (https://github.com/r3fang/snATAC) with slight modification. For the data in each developmental stage, pair-end sequencing reads were aligned to mouse genome (mm10) using bowtie2, non-uniquely mapped and improperly paired alignments were filtered, PCR duplications and mitochondrial reads were removed. Macs2 software was used to perform peak calling on retained reads and a read count matrix was generated with peaks in the row and cells in the column. The read count matrix in peaks was then converted to the matrix in promoters, by merging the peaks located in same promoter. The promoter read count matrix was used to perform dimension reduction analysis to cluster and assign the cells into known cell types. Cells with reads located in promoter of *Neurod6* were defined as excitatory neurons, cells with reads located in promoter of *Dlx5* and without reads located in promoter of *Hes5* were defined as inhibitory neurons [29]. Finally, the excitatory and inhibitory neurons were merged, respectively, to produce a pseudo-bulk ATAC-seq dataset for each neuronal subtype in each developmental stage.

### Analysis of scRNA-seq data

scRNA-seq data from developing mouse brain at E12.5, E13.5, E14.5, E15.5, E16.5, P0, P7, P21 and P60 were downloaded from previous studies [31–33] (**Table S2**), a read count matrix with gene in row and cell in column was achieved in each stage. Seurat [51] was adopted to perform data analysis. Briefly, the excitatory and inhibitory neurons were selected and retained for downstream analysis according to the annotations in raw datasets. Remaining neurons with potential double droplets or having mitochondrial mRNA loads over 10% were removed. The genes expressed in less than 3 cells and the cells expressed less than 200 genes were filtered in further analysis. The retained expression read count matrix was normalized by NormalizeData function, and top 2000 variable genes were selected from the normalized matrix using FindVariableFeatures function. Dimensional reduction was performed based on normalized expression matrix of top 2000 variable genes, and the top 10 principal components were used to generate the UMAP (Uniform Manifold Approximation and Projection).

### Integration of scRNA-seq and snATAC-seq datasets

The Seurat pipeline was adopted to integrate scRNA-seq and snATAC-seq datasets for each development stage. Briefly, gene expression matrix (scRNA-seq data) and promoter coverage matrix (snATAC-seq data) were normalized and scaled by using NormalizeData function and ScaleData function, separately. FindTransferAnchors function was used to find anchors (shared genes) between scRNA-seq and snATAC-seq datasets. The scaled gene expression matrix and promoter coverage matrix were transferred and merged into a co-embedded matrix based on the anchors by using canonical correlation space (CCA) analysis. The co-embedded matrix was scaled *via* ScaleData function, and dimensional reduction was performed on scaled co-embedded matrix.

### Expression clustering analysis

Clustering analysis for EXC- and INH-predominant EGR1 binding sites associated genes was performed by Mfuzz software [52]. By default, a fuzzy c-means clustering was performed on expression matrix with gene in row and sample in column. The fuzzifier parameter (*m*) was estimated by mestimate function in the Mfuzz. The number of clusters (*c*) was set to 5, since it is the minimum cluster number satisfies the condition that one of the clusters showing similar expression pattern with *Egr1*.

## Authors’ contributions

H. X. conceived and designed the study; H. X., M. F., and X. L. supervised the study; B. C., G. C. and M. F. provided and characterized mouse strains; X. X. isolated neurons and constructed CUT&RUN libraries; L. Y., Y. L. and Y. C. performed the bioinformatic analyses; L. Y., X. X., and H. X. interpreted results and wrote the manuscript. All authors discussed the results and edited the manuscript.

## Availability of data and materials

The datasets supporting the conclusions of this article are available in the NCBI Gene Expression Omnibus (GEO) with the accession number GSE218312. Publicly available datasets used in this study are summarized in Table S2. All the other data generated in this study are included in the article and the additional files. Data analysis scripts used in this study are available on GitHub repository (https://github.com/Gavin-Yinld/Neuronal_CUT.RUN).

## Supplementary information

Additional file 1: Supplemental figures 1-8.

Additional file 2: Supplemental table 1. Reproducible peaks between biological replicates for histone modifications of excitatory and inhibitory neurons.

Additional file 3: Supplemental table 2. A summary of public datasets used in this study.

Additional file 4: Supplemental table 3. Super enhancers of excitatory and inhibitory neurons. Additional file 5: Supplemental table 4. Reproducible EGR1 peaks between biological replicates for excitatory and inhibitory neurons.

