## Supplemental Figures for "Elevated EGR1 Binding at Enhancers in Excitatory Neurons Correlates with Neuronal Subtype-Specific Epigenetic Regulation"

### Figure S1

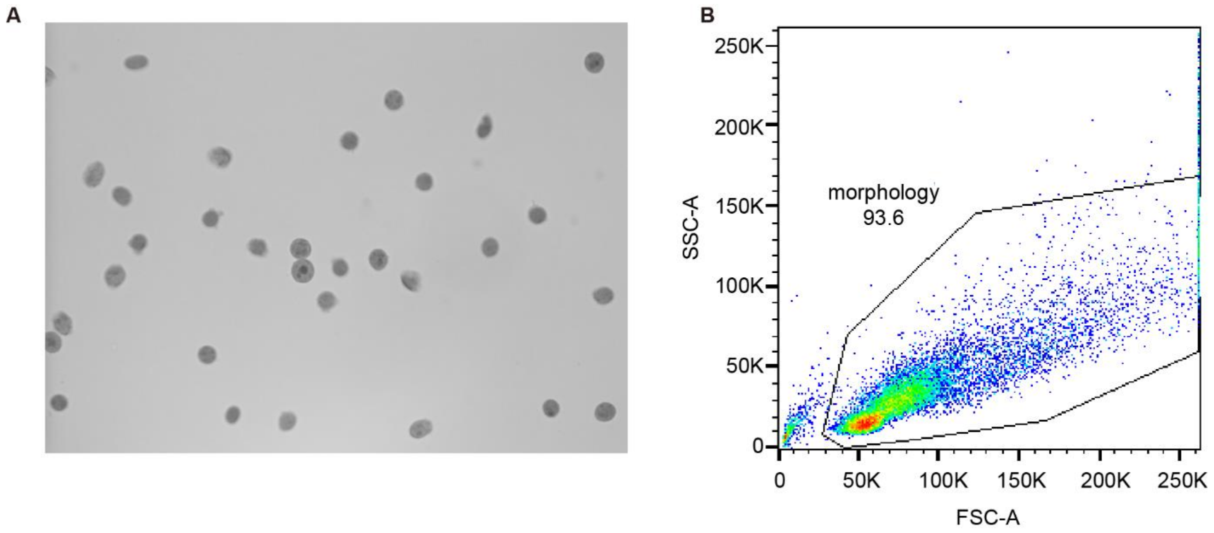


**Figure S1**. **Removal of cell debris. A)** The image of single nuclei suspension preparation was taken by microscope. **B)** Flow cytometry showed the majority of the suspension were nuclei and it showed clear separation between the nuclei and the cell debris.

### Figure S2

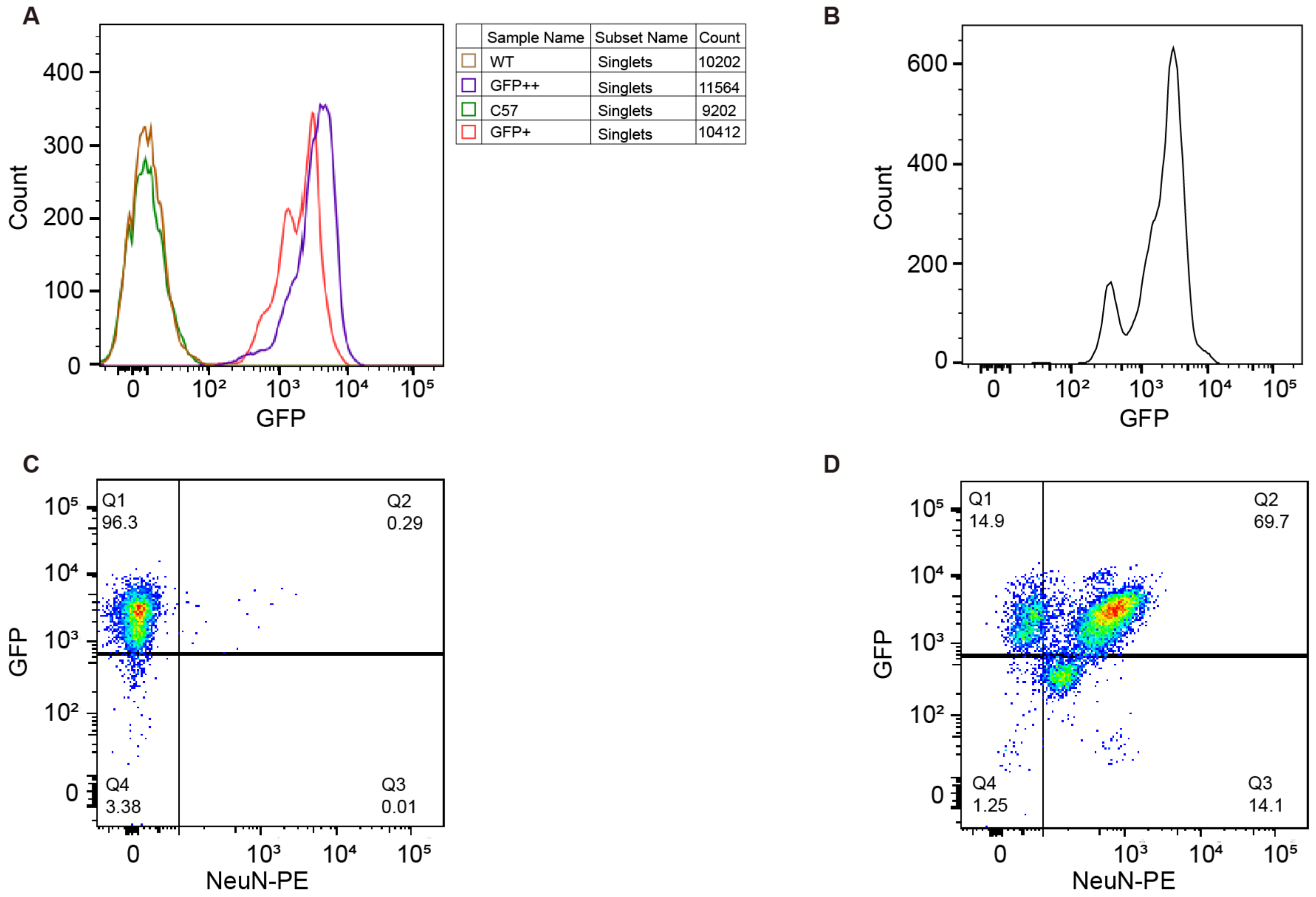


**Figure S2**. **FACS-sorting control and gating. A)** Signal intensity of GFP in WT mouse brain samples and Sun1 flox | Emx1-Cre (+) mouse brain samples. **B)** Signal intensity of GFP in a representative Sun1 flox | Emx1-Cre (+) mouse brain sample. **C)** FACS-sorting for Sun1 flox | Emx1-Cre (+) mouse brain sample (control sample) without anti-NeuN-PE Ab staining. **D)** FACS-sorting for Sun1 flox | Emx1-Cre (+) mouse brain sample with anti-NeuN-PE Ab staining.

### Figure S3

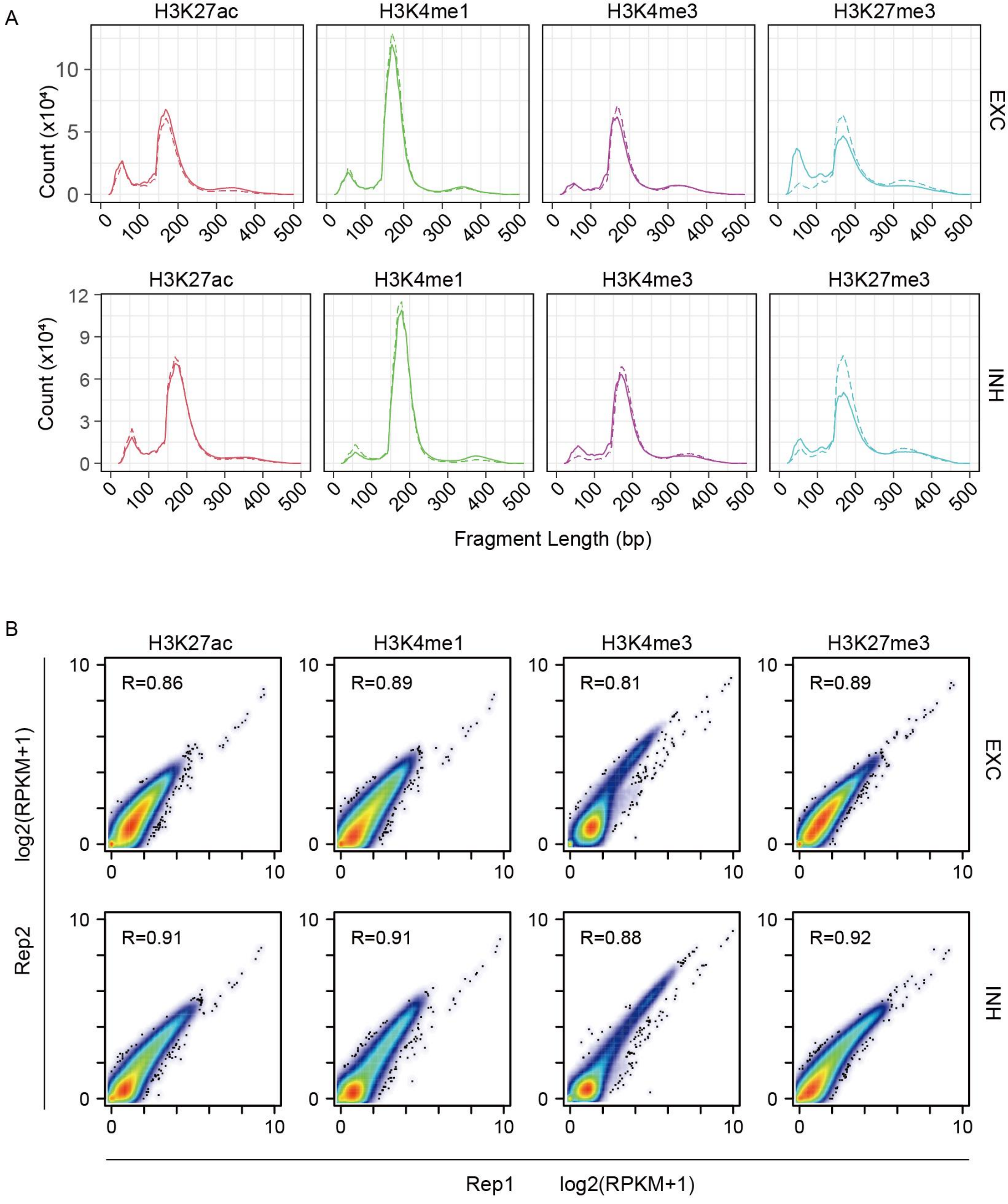


**Figure S3. Confirmation of high quality CUT&RUN libraries. A)** Fragment size distribution for each sample. The dashed and solid lines represent two biological replicates, respectively. **B)** Correlation between biological replicates for each histone modification in excitatory and inhibitory neurons.

### Figure S4

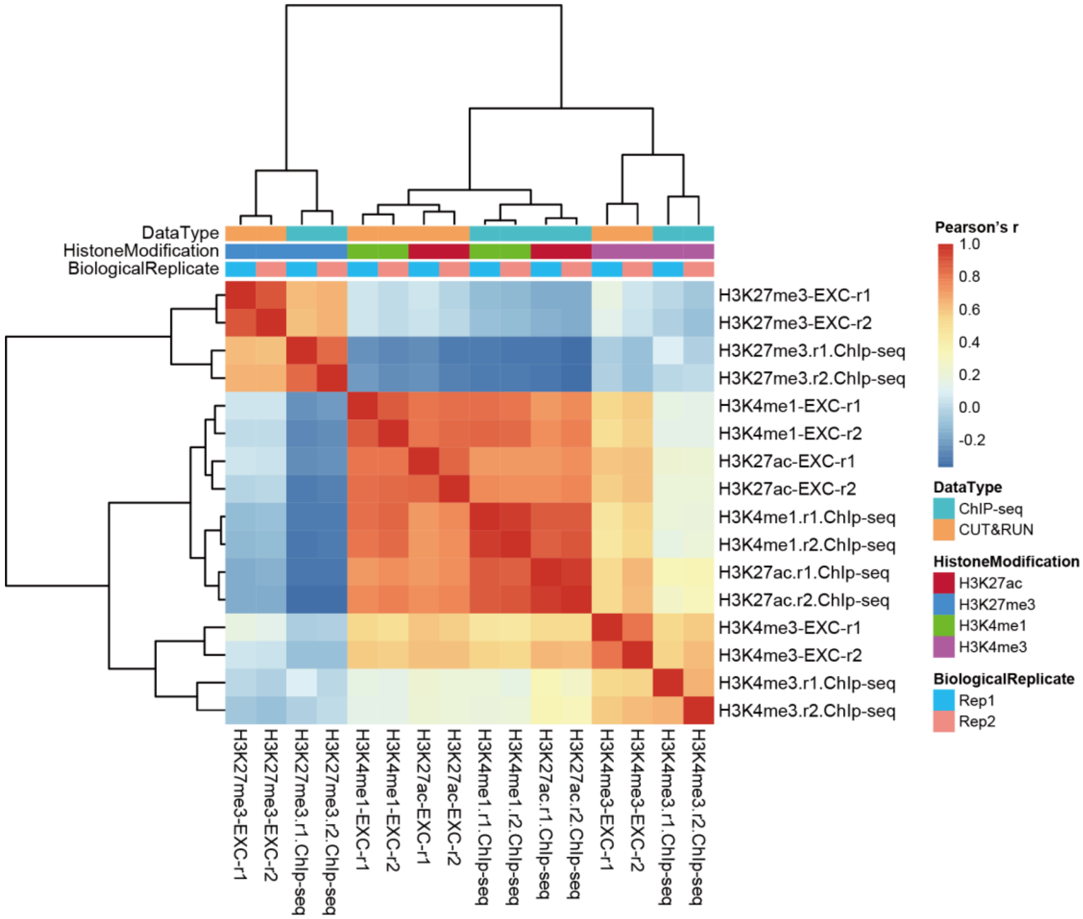


**Figure S4. Correlation between CUT&RUN data in this study and ChIP-seq data from (Mo et al., Neuron, 2015) for excitatory neurons.**

### Figure S5

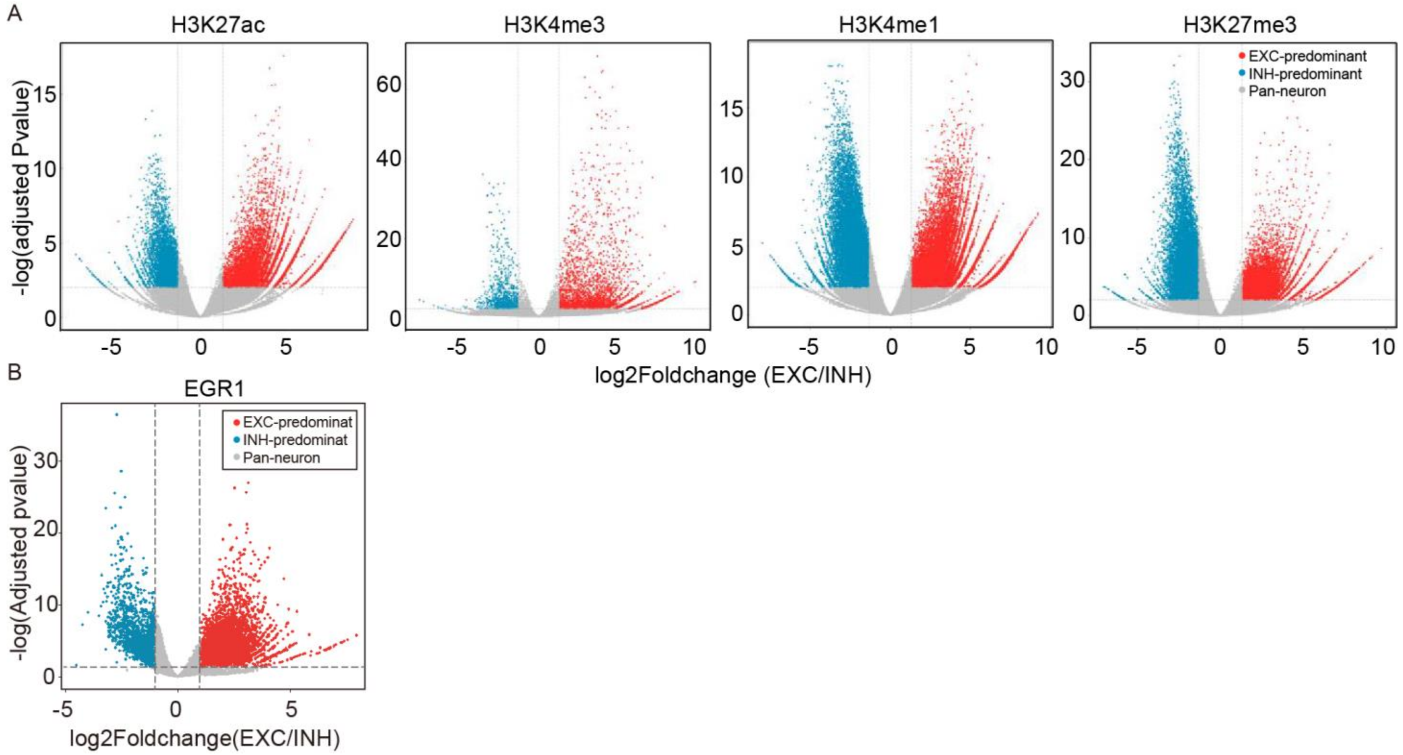


**Figure S5. Comparison of histone modifications (A) and EGR1 binding (B) between excitatory and inhibitory neurons.** The differential peak regions colored in blue and red were determined by “result” function in the DESeq2, with thresholds “padj≤0.05” and “FoldChange≥2” or “FoldChange≤0.5” (see Methods).

### Figure S6

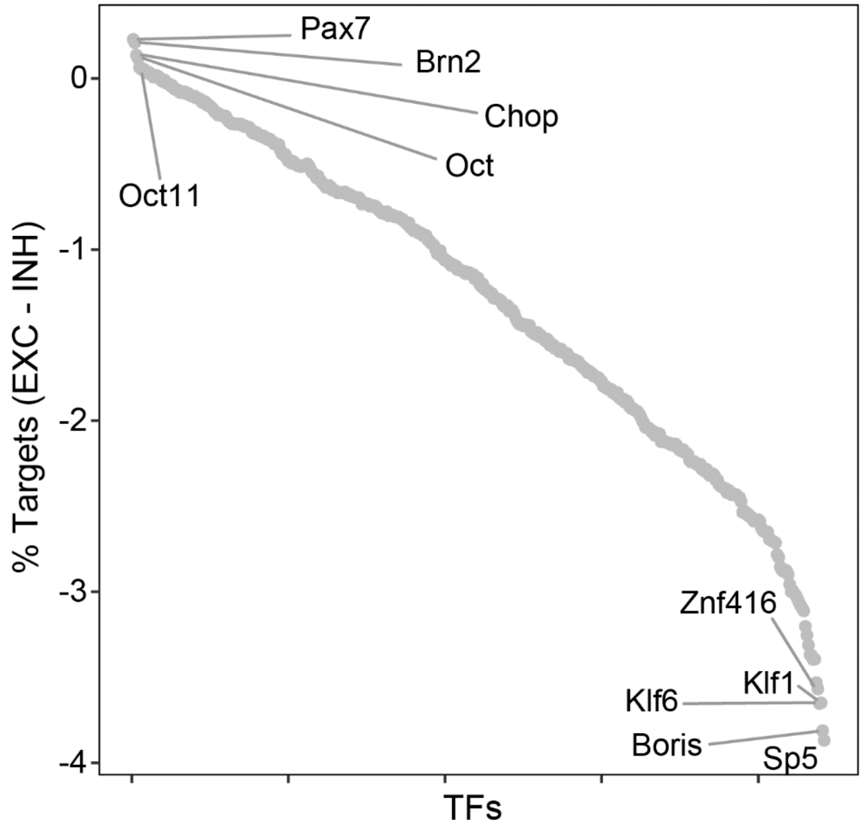


**Figure S6. Scatterplot showing the proportion changes of TFs' motif at active promoters of excitatory neurons compared with inhibitory neurons.** For each motif, the percentage of active promoters containing this motif was calculated for excitatory and inhibitory neurons, separately. The promoter containing difference for each motif in two neuronal subtypes is shown in y-axis.

### Figure S7

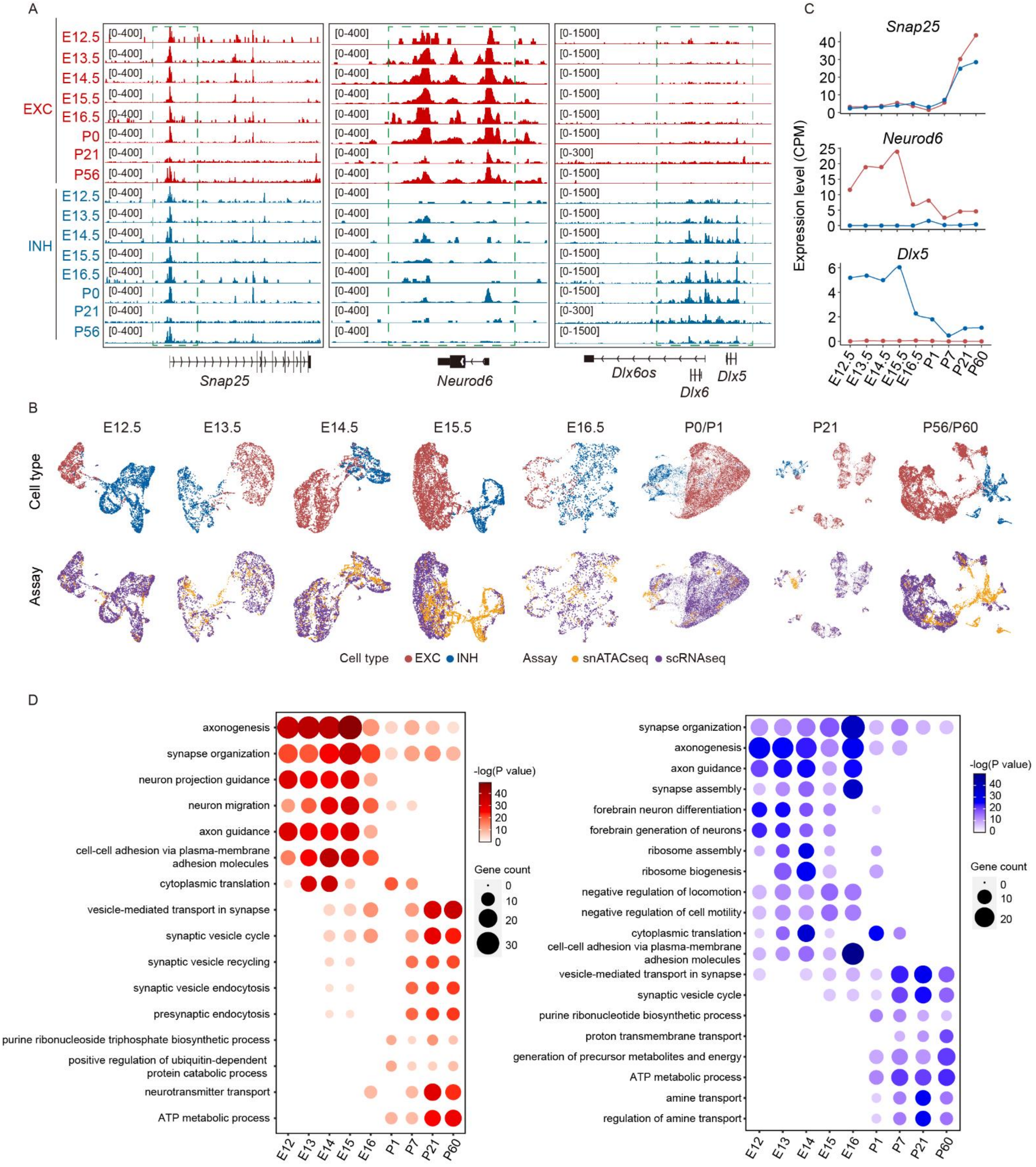


**Figure S7. Integration of scRNA-seq and snATAC-seq data during mouse brain development. A)** Examples to show chromatin accessibility surrounding neuronal marker genes during brain development. **B)** Integration of scRNA-seq and snATAC-seq datasets in each developmental stage. **C)** Expression of neuronal marker genes related to A). **D)** Function of top 200 stage-specific expressed genes in each developmental stage. Gene specificity was defined as $CPM_{i,t}/median(CPM_{(i,\ all\ stages)})$, where i represents for gene and t represents for stage.

### Figure S8

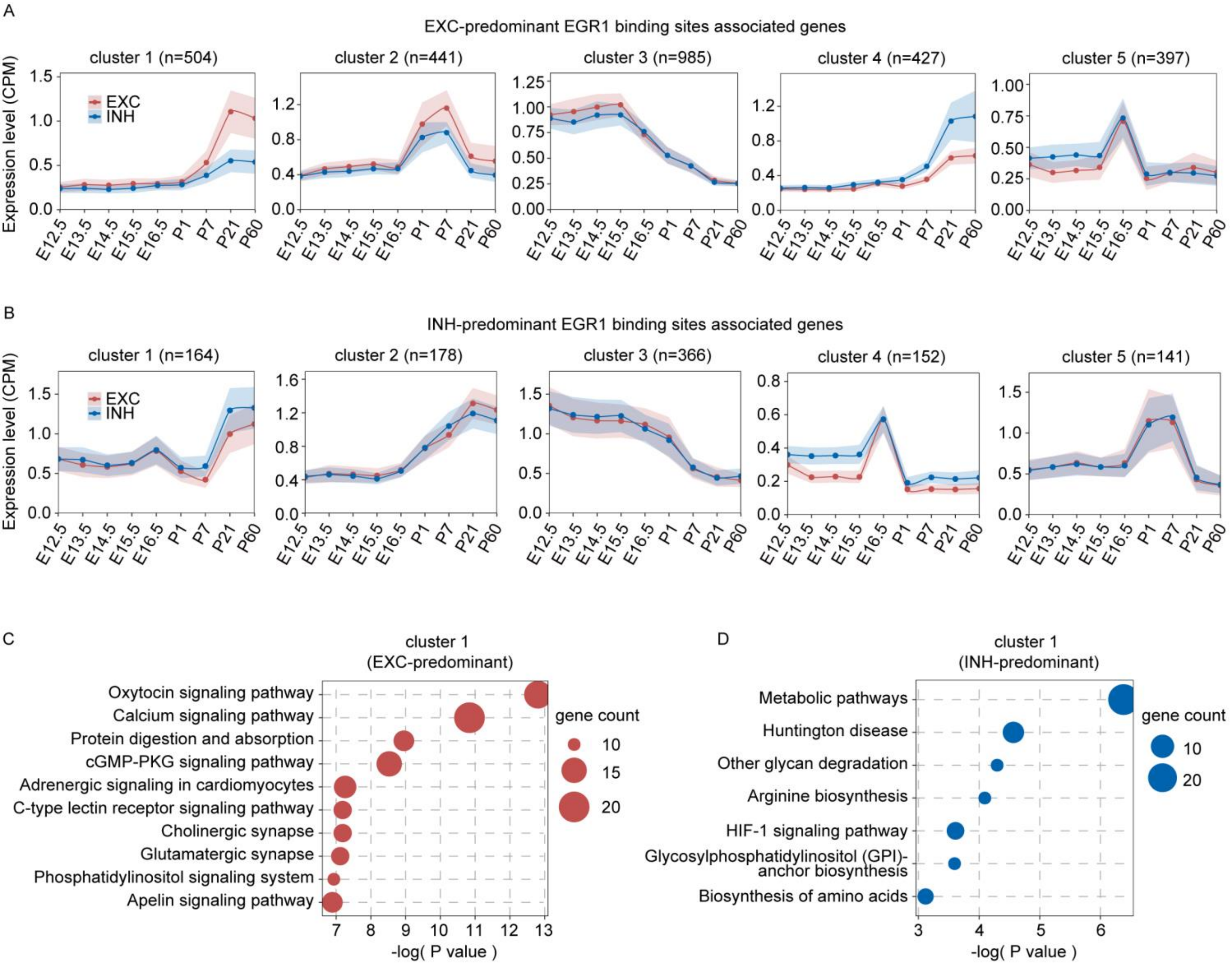


**Figure S8. Expression clustering and gene ontology analysis for EXC- and INH-predominant EGR1 peaks associated genes. A and B)** Expression clustering analysis for EXC- (A) and INH-predominant (B) EGR1 peaks associated genes. **C)** KEGG pathway enrichment analysis for cluster 1 genes among all EXC-predominant EGR1 peaks associated genes. **D)** KEGG pathway enrichment analysis for cluster 1 genes among all INH-predominant EGR1 peaks associated genes.
